# AptCancerDB: A Curated Knowledgebase and Translational Discovery Platform for Anticancer Aptamers

**DOI:** 10.64898/2026.07.02.735999

**Authors:** Nisha Bajiya, Shivansh Singh, Gajendra P. S. Raghava

## Abstract

Aptamers are emerging as important molecular recognition ligands in oncology, playing significant roles in cancer diagnostics, targeted therapies, drug delivery systems, and molecular imaging. Numerous aptamers have advanced to clinical trials, indicating their potential for real-world applications; however, existing databases fail to capture that. To bridge this critical gap, we developed AptCancerDB (https://webs.iiitd.edu.in/raghava/aptcancerdb/), a comprehensive, manually curated database of experimentally verified anticancer aptamers. The current release contains 1,941 entries collected from studies published between 2000 and 2025, covering 29 cancer types, approximately 200 cancer cell lines, and direct links to 22 clinical trials. Each entry is annotated with sequence information, target details, cancer type, cell line, SELEX methodology, affinity determination data, chemical modifications, and biological activities. The dataset is dominated by 82.7% ssDNA, reflecting its superior stability and ease of synthesis, while only 16.6% is ssRNA and appears primarily in studies targeting complex intracellular or protein-protein interactions. To facilitate structural analysis, predicted secondary structures, dot-bracket notations, specific structural elements, and minimum free energy values were also included. AptCancerDB integrates a MySQL backend with an ArcadeDB/OpenCypher-based Knowledge Graph, enabling exploration of relationships among aptamers, targets, cancer types, cell lines, and functional applications. The platform provides advanced search and browsing facilities, BLASTn-based similarity searching, and GC Calculator. Built on a modern, responsive frontend (React/TypeScript/Tailwind CSS), the platform includes a REST API for data retrieval. By integrating fragmented experimental data into a unified cancer-focused resource, AptCancerDB serves as a valuable resource for comparative analysis, aptamer discovery, and the development of next-generation aptamer-based diagnostics and therapeutics.

**Highlights:**

- Curated knowledge base of experimentally validated anticancer aptamers.
- AptCancerDB contain therapeutic, tumor-homing and cell-penetrating aptamers.
- Summarizes clinical progress and translational trends in anticancer aptamer research.
- Supports rational aptamer design using molecular, functional, and clinical annotations
- Disease-focused resource for cancer diagnosis, therapy, and drug delivery

**Teaser:** AptCancerDB maintains experimentally validated anticancer aptamers relevant to diagnosis, drug delivery, and therapy.

## 1. Introduction

Cancer remains a leading cause of global morbidity and mortality and continues to pose major therapeutic challenges due to tumor heterogeneity, adaptive resistance, and the toxicity of conventional treatments (1). In 2022, approximately 20 million new cancer cases and 9.7 million deaths were reported worldwide, with projections exceeding 35 million new cases by 2050 [<u>WHO</u>]. Consistent with this burden, cancer is a major cause of mortality among men aged 60-79 years and women aged 40-79 years (2). Although surgery, radiotherapy, and cytotoxic chemotherapy remain central to cancer treatment, their clinical utility is frequently limited by systemic toxicity, poor tumor selectivity, and the emergence of drug resistance (3). The side effects of chemotherapy and organ toxicity highlight the need for more precise treatment strategies (4,5). Targeted therapies and immunotherapies have improved outcomes in certain cases, but existing therapies face a number of challenges, including limited tumor penetration, immunogenicity, complex manufacturing, and off-target effects. Hence, there is a need for alternative molecular recognition agents to facilitate selective targeting and improve clinical outcomes (6,7).

To address these challenges, aptamers are gaining interest as a promising class of molecular recognition ligand for cancer treatment. Aptamers are short, single-stranded (ss) nucleic acids, typically DNA or RNA oligonucleotides, that exhibit strong affinity and high specificity for their target molecules (8). Unlike linear oligonucleotides, aptamers fold into distinct secondary and tertiary structures, such as stems, loops, hairpins, and G-quadruplexes. This unique folding allows them to adapt and change shape to recognize targets effectively (9). Target binding occurs through various noncovalent interactions, including hydrogen bonding, π–π stacking, van der Waals forces, and electrostatic interactions. Due to this structural flexibility and range of interactions, aptamers can accurately distinguish closely related targets, making them highly attractive ligands for selective cancer targeting and translational applications (10).

The tumor-homing and delivery capabilities of aptamers are enabled by robust selection technologies. Aptamer discovery began in 1990 with the development of the SELEX (Systematic Evolution of Ligands by Exponential Enrichment) method for the in vitro evolution of nucleic acid ligands (11). In parallel, Ellington and Szostak demonstrated the directed evolution of oligonucleotides capable of precise molecular recognition, introducing the term “aptamer,” derived from the Latin *“aptus”* (meaning “to fit”) and the Greek “meros” (meaning “region”) (12). SELEX allows the isolation of functional aptamers from large random single-stranded nucleic acid libraries through iterative rounds of binding, partitioning, and amplification (10). Over time, SELEX has evolved into several specialized formats, including cell-SELEX, in vivo SELEX, capture-SELEX, microfluidic SELEX, and high-throughput sequencing-assisted SELEX, enabling selection against purified proteins, intact cells, tissues, and heterogeneous tumor phenotypes (13,14). These advances have facilitated the generation of cancer-selective aptamers with binding affinities frequently reaching the low-nanomolar to picomolar range, supporting their application in targeted therapy, delivery, and diagnostics (15).

Aptamers exhibit antibody-like specificity due to their ability to adopt complex three-dimensional structures and are therefore often referred to as “chemical antibodies” (16). Aptamers offer several advantages over protein-based ligands, such as monoclonal antibodies, particularly applicable in translational oncology. They can be chemically synthesized with high reproducibility, have low immunogenicity, and enable targeted modifications to improve stability and pharmacokinetics (14)(17–21). Therapeutic aptamers function as antagonists that tend to inhibit protein-protein interactions via directly binding to tumor-associated receptors or ligands (e.g., EGFR, VEGF, CXCL12) (22–24). It prevents their interaction with natural partners, thereby inhibiting oncogenic signaling pathways, reducing cell motility and invasion, and inducing apoptosis. This ultimately suppresses cancer cell proliferation, migration, and survival, similar to the activity of anticancer peptides (ACPs) (25). Their small size facilitates efficient tumor penetration, enabling selective accumulation at tumor sites. In this context, many anticancer aptamers function as tumor-homing ligands, analogous to tumor-homing peptides, by selectively recognizing cancer cells or tumor-associated targets (26,27).

Advances in aptamer-guided drug delivery have substantially improved targeting precision in cancer treatment. In this, the tumor-homing property has been extensively exploited for targeted drug delivery, including aptamer-drug conjugates (ApDCs) and aptamer-functionalized nanocarriers, such as nanoparticles, liposomes, and micelles. These platforms enable receptor-mediated internalization, enhance drug accumulation within tumors, and reduce off-target toxicity, thereby strengthening the translational potential of aptamer-based cancer therapeutics and diagnostics (28).

Beyond delivery, aptamers are being explored to overcome drug resistance by targeting multidrug resistance transporters, modulating anti-apoptotic signaling, or delivering siRNA and antisense oligonucleotides that silence resistance-associated genes (29). Importantly, a subset of aptamers exhibits cell-penetrating properties, enabling efficient cellular uptake and intracellular trafficking through multiple mechanisms, analogous to cell-penetrating peptides, and thereby expanding their utility for intracellular targeting (30,31). Aptamer-based biosensors and imaging probes, such as PET, MRI, and fluorescence platforms, facilitate early detection and tumor-specific imaging (32–37). Also, liquid biopsy techniques that use aptamers can detect circulating tumor cells and biomarkers with minimal invasiveness (38,39). Additionally, integrating aptamer-based treatments with chemotherapy or immunotherapy may enhance the overall effectiveness of the treatment (40,41). Together, these developments depict aptamers as versatile molecular tools that play a crucial role in precision oncology.

Aptamer-based technologies have significant potential but face several translational challenges. Key limitations include rapid renal clearance and are vulnerable to breaking down due to nucleases, which require chemical stabilization or carrier strategies. After cellular uptake, endosomal trapping often hinders the efficient cytosolic delivery of therapeutic payloads. Moreover, identifying and verifying appropriate cell-surface biomarkers is difficult due to variation among tumor subclones and different patient populations. These issues underscore the need for improved selection pipelines, reliable in vivo validation models, and integrated computational methods to predict aptamer-target interactions, stability, and pharmacokinetics (15). Addressing these challenges necessitates the systematic integration of scattered experimental data, standardized annotation of aptamer properties, and creating resources focused on diseases for comparative analysis and rational design, leading to the development of comprehensive platforms like AptCancerDB.

### 1.1. Anticancer aptamers in Clinical trials

The translational advancement of aptamer technology has now reached the clinical stage, emphasising its potential as both therapeutic and diagnostic agents in oncology. A total of 22 aptamer-associated clinical trials (https://clinicaltrials.gov/) have been registered across various cancer types, including solid tumours and haematological malignancies, as well as pancreatic, renal, lung, and colorectal cancers. Several anticancer aptamer-based candidates have entered Phase I and II clinical trials, demonstrating acceptable safety profiles, tumor-specific accumulation, and promising pharmacodynamic outcomes. As summarized in Table 1, the current registration involving anticancer aptamers includes a mix of completed, ongoing, and recruiting trials. Out of these, approximately 9 have been completed, 9 are in unknown or pending status, 2 are actively recruiting, and 2 have been withdrawn or terminated.

**Table 1:**
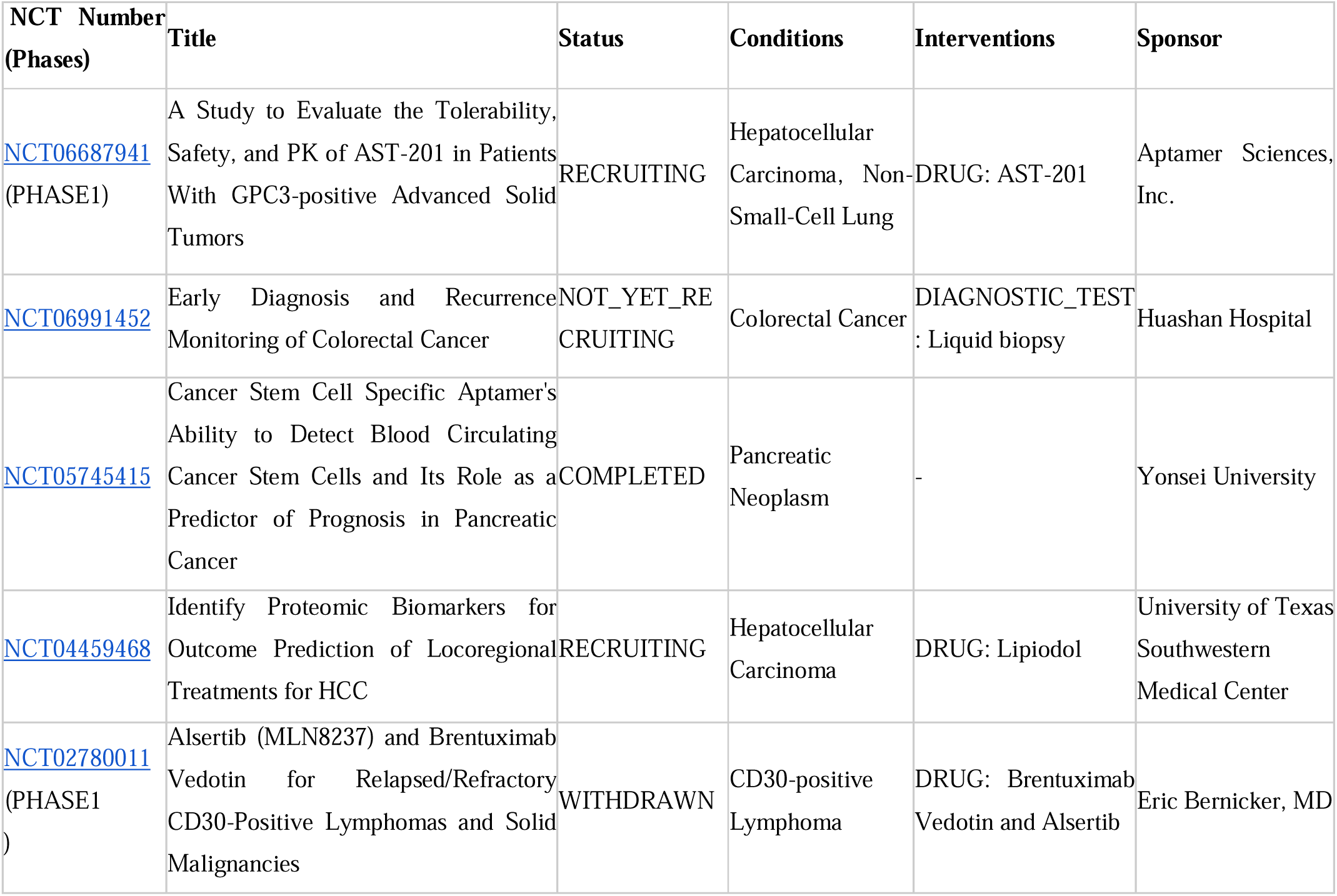
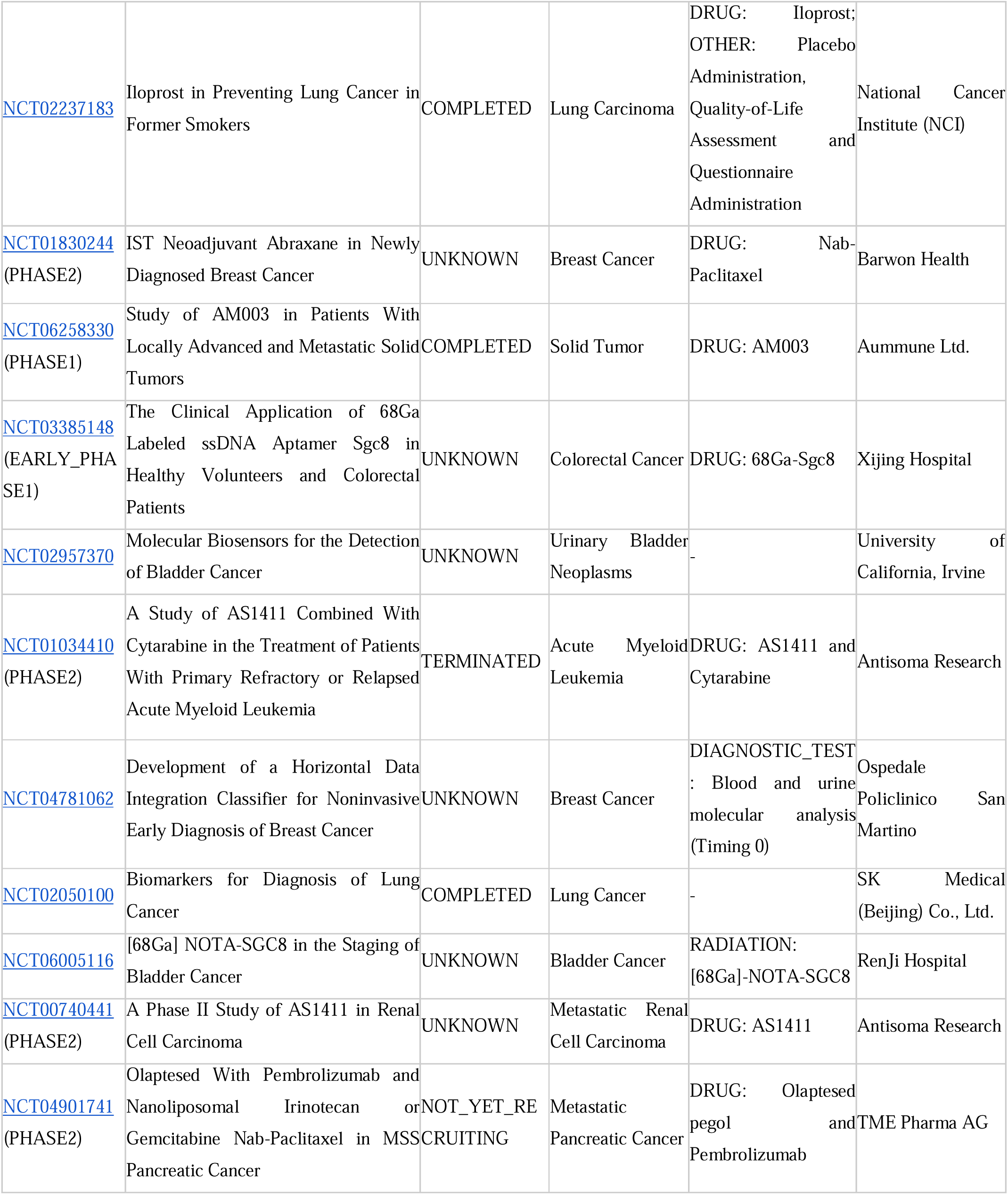
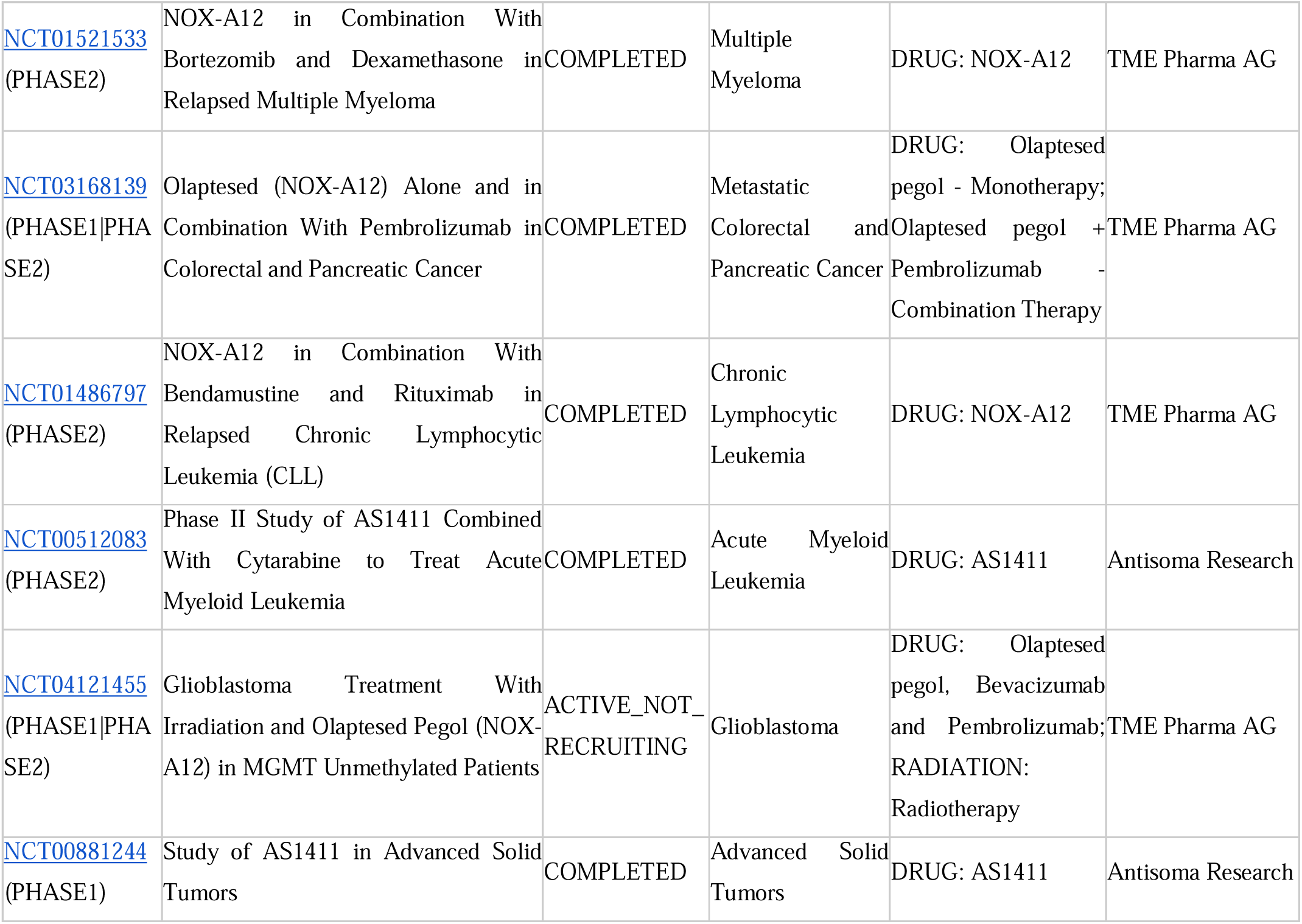
List of Clinical trials of aptamers in Cancer-related studies.

In oncology, the most advanced candidates in development are AS1411 and NOX-A12 (Olaptesed pegol). AS1411, a G-quadruplex-forming oligodeoxynucleotide and one of the first and most extensively studied DNA aptamers to enter clinical evaluation, targets nucleolin and has been assessed across various cancer types, including leukemia and renal carcinoma, validating its safety and tumor-targeting capability. Similarly, another promising aptamer, NOX-A12 (Olaptesed pegol), a PEGylated Spiegelmer-type aptamer targeting CXCL12, inhibits interaction with its receptors, CXCR4/CXCR7, and has advanced through several early- and mid-phase trials, including combinatorial regimens with immune checkpoint inhibitors such as pembrolizumab, demonstrating synergistic efficacy and a manageable toxicity profile.

Beyond therapeutics, diagnostic and imaging applications of aptamers are also emerging in clinical settings. Examples include 68Ga-labelled ssDNA aptamer Sgc8, designed for PET-based imaging in colorectal and bladder cancers, and liquid biopsy platforms employing aptamers for early detection of circulating tumor cells or tumor-derived nucleic acids. These studies underscore the versatility of aptamers as both therapeutic agents and diagnostic biosensors.

### 1.2. Existing Aptamer Databases

Traditional aptamer selection relies on the SELEX process, in which an extensive random oligonucleotide library (∼10^15^ sequences) is iteratively incubated with a target, separating bound from unbound molecules, followed by PCR amplification of the enriched pool (42). Multiple cycles yield high-affinity, high-specificity aptamers that are subsequently sequenced and validated using binding assays. Although highly effective, SELEX is labor-intensive, often requiring weeks to months, and the initial library composition strongly influences its output. As a result, potentially valuable sequences may be lost during the selection process, thereby compromising overall discovery efficiency (43). To address these issues, researchers are utilising artificial intelligence (AI) methods to facilitate the rapid design and discovery of new aptamers (44).

Over the past two decades, several aptamer-specific resources have been developed, including sequence, structural, and experimental information generated from SELEX and related methodologies. The first dedicated aptamer repository was established in 2004 by Lee et al. (45) and later updated in 2024 as the UTexas Aptamer Database (46). Since then, the landscape has been expanded by additional resources, including AptaDB, which contains experimentally validated aptamer-target interaction data (47). Other additional nucleic acid-based aptamer databases have been reported in Table 2.

**Table 2.**
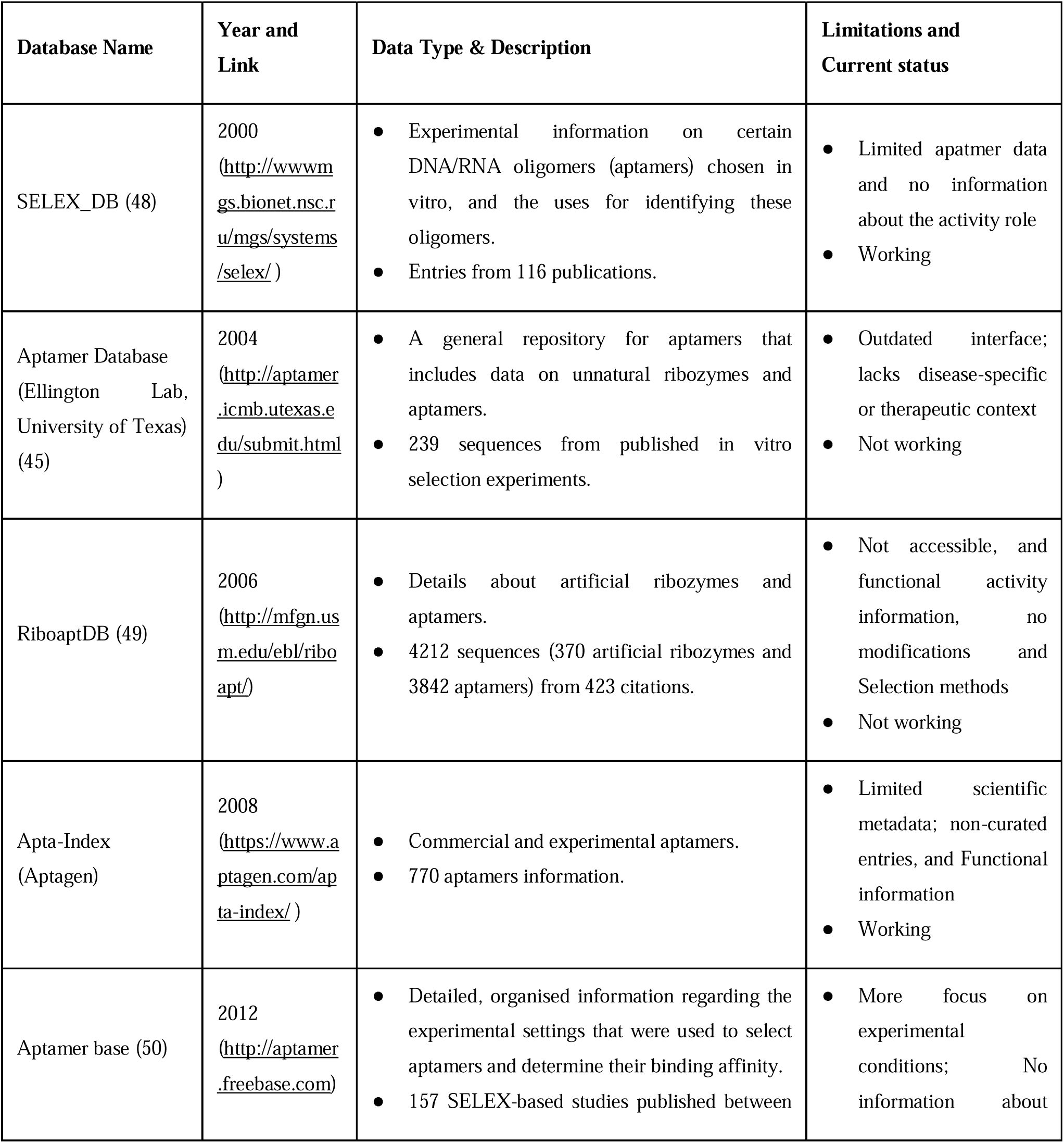
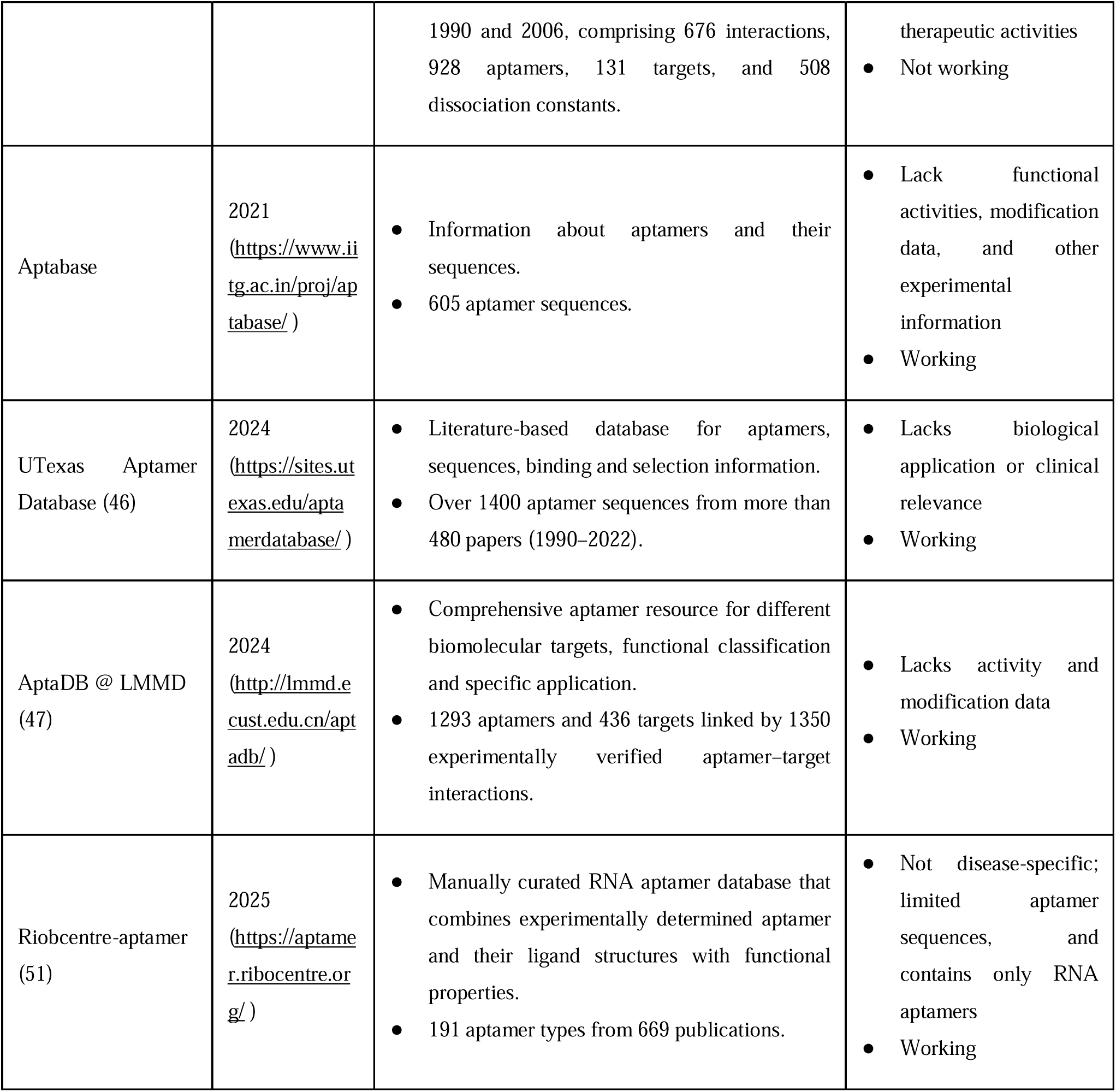
Comparison of Existing Aptamer Databases developed over the years.

### 1.3. Need for AptCancerDB

Significant advances have been made in developing computational resources for cancer therapeutics, with efforts predominantly focused on ACPs. Over the past two decades, numerous ACP databases such as CancerDR (52), ApInAPDB (53), CancerPPD2 (54) and APD6 (55), together with several machine-learning-based prediction tools, have greatly promoted peptide-based cancer therapeutics and biomarker discovery (56–59). These resources aid in a comprehensive understanding of peptide sequences, biological activities, physicochemical properties, and predictive models that support rational therapeutic design. However, despite advances in peptide-based cancer bioinformatics, existing aptamer repositories remain limited in scope and lack sufficient biochemical data (sequences, targets, Kd values, and SELEX conditions), as well as contextual annotation needed for translational oncology research (51).

Therefore, unlike anticancer peptides, there is currently no publicly available resource that focuses exclusively on experimentally validated anticancer aptamers with comprehensive functional and translational annotations. This is particularly constrained considering the ever-expanding range of aptamer research directed toward cancer, including aptamers selected against tumour-specific receptors and biomarkers, cancer-cell binders from cell-SELEX, aptamer-drug and aptamer-siRNA conjugates, immunomodulatory aptamers, and a variety of diagnostic and theranostic applications (29)(39)(60,61). Hence, there is an urgent need for a centralised, well-curated platform that compiles cancer-specific aptamer sequences, target annotations, chemical modifications, biological roles and experimental outcomes to enable rational design, accelerate discovery and bridge the gap between preclinical research and clinical translation in oncology.

To address this unmet need, we present AptCancerDB, an extensive, thoroughly maintained resource dedicated solely to experimentally verified aptamers for cancer therapy, diagnosis and research. AptCancerDB is a disease-focused resource that combines multidimensional metadata such as screening and selection details, sequence and structural features, target cancer type, target biomolecule, cell line used, SELEX strategy, affinity determination method, chemical modifications, application category, and, when available, inhibitory outcomes, clinical-trial information, and patent data. Besides supporting rational sequence and motif analysis for next-generation aptamer engineering, its design offers a translational link between basic research and real-world clinical applications. Overall, AptCancerDB serves as a comprehensive analytical tool designed for cancer research, providing researchers with a focused, data-driven space to explore, compare, and develop aptamer-based diagnostics and treatments.

## 2. Materials and Methods

### 2.2. Data Collection Strategy

AptCancerDB was developed through an extensive, multi-step process for data collection and curation, combining existing aptamer databases with significant mining of published literature. Initially, data were retrieved from well-known public databases, such as the UTexas Aptamer Database (46), AptaBase (www.iitg.ac.in/proj/aptabase), and AptaDB (47). Duplicate articles were removed, retaining only unique ones from all three databases. Following this, a systematic literature search was conducted across PubMed and Google Scholar (https://scholar.google.com.hk/) to identify additional studies on cancer-related aptamers. The search strategy employed combinations of relevant keywords, including ‘Aptamers’, ‘Antitumor/Anticancer aptamers’, ‘Therapeutic aptamers’, and ‘Diagnostic aptamers’. Publications from the last 25 years (2000-2025) were reviewed to provide a comprehensive overview of the most recent advancements in anticancer aptamer research. This combined search yielded 950 PubMed-indexed papers and 36 DOI of research articles. All selected publications were thoroughly reviewed, and manually curated data on aptamers used in cancer diagnosis, treatment, and targeted delivery were compiled. After eliminating identical articles and incomplete records, the final curated dataset comprised 1,941 aptamer entries. Before being included in our resource, every entry was carefully examined to ensure its accuracy and quality. The complete database framework is shown in Figure 1.

**Figure 1:**
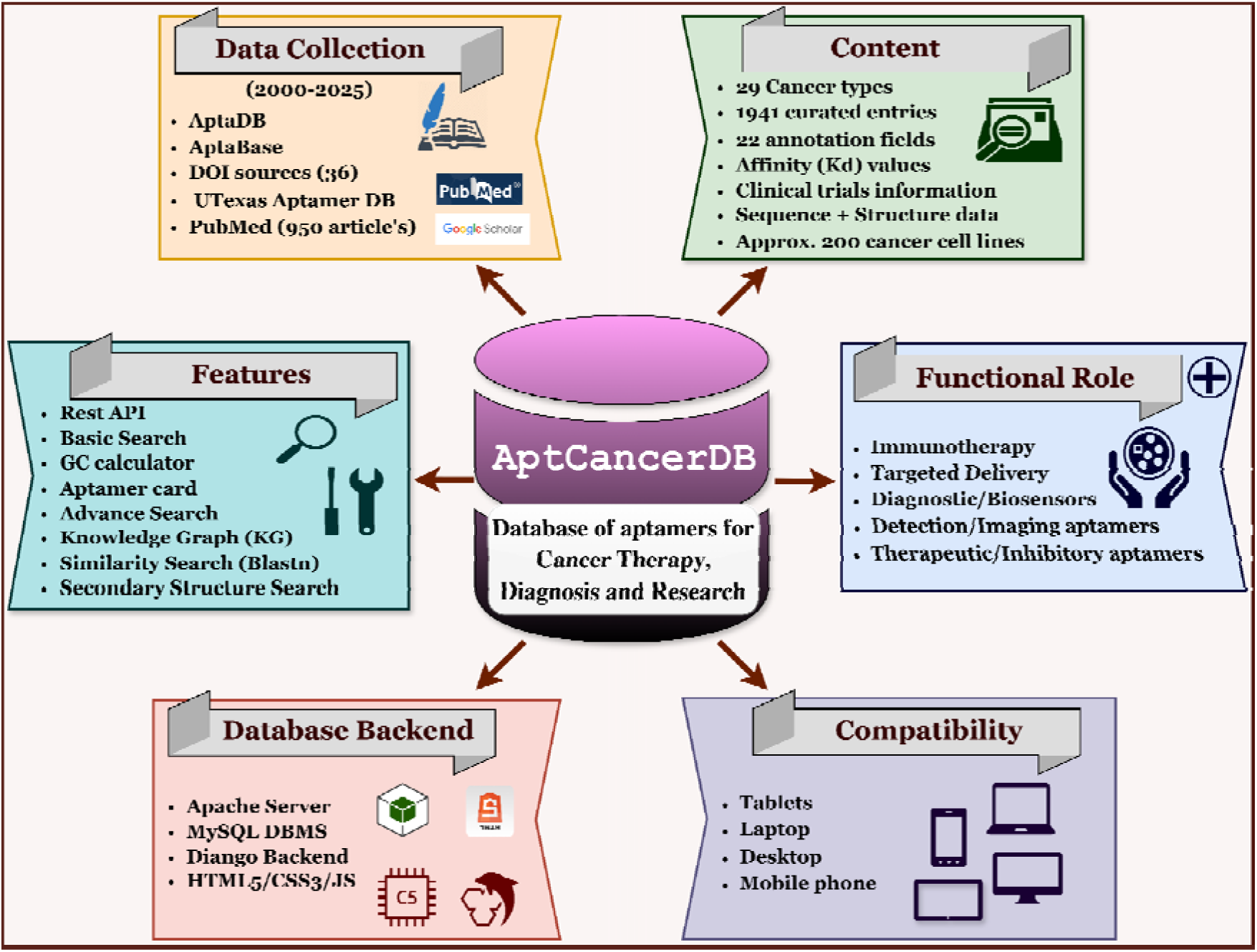
The Architecture of the AptCancerDB database.

### 2.2. Data Annotation

AptCancerDB is a carefully organized collection of anticancer aptamer sequences, lengths, experimental methods, and inhibitory effects. Table 3 presents the annotations for the 22 key fields that comprise this database. Each entry links directly to either the PubMed ID (PMID) or DOI of the article, allowing readers to easily access the original research journal from which the information was retrieved.

**Table 3:**
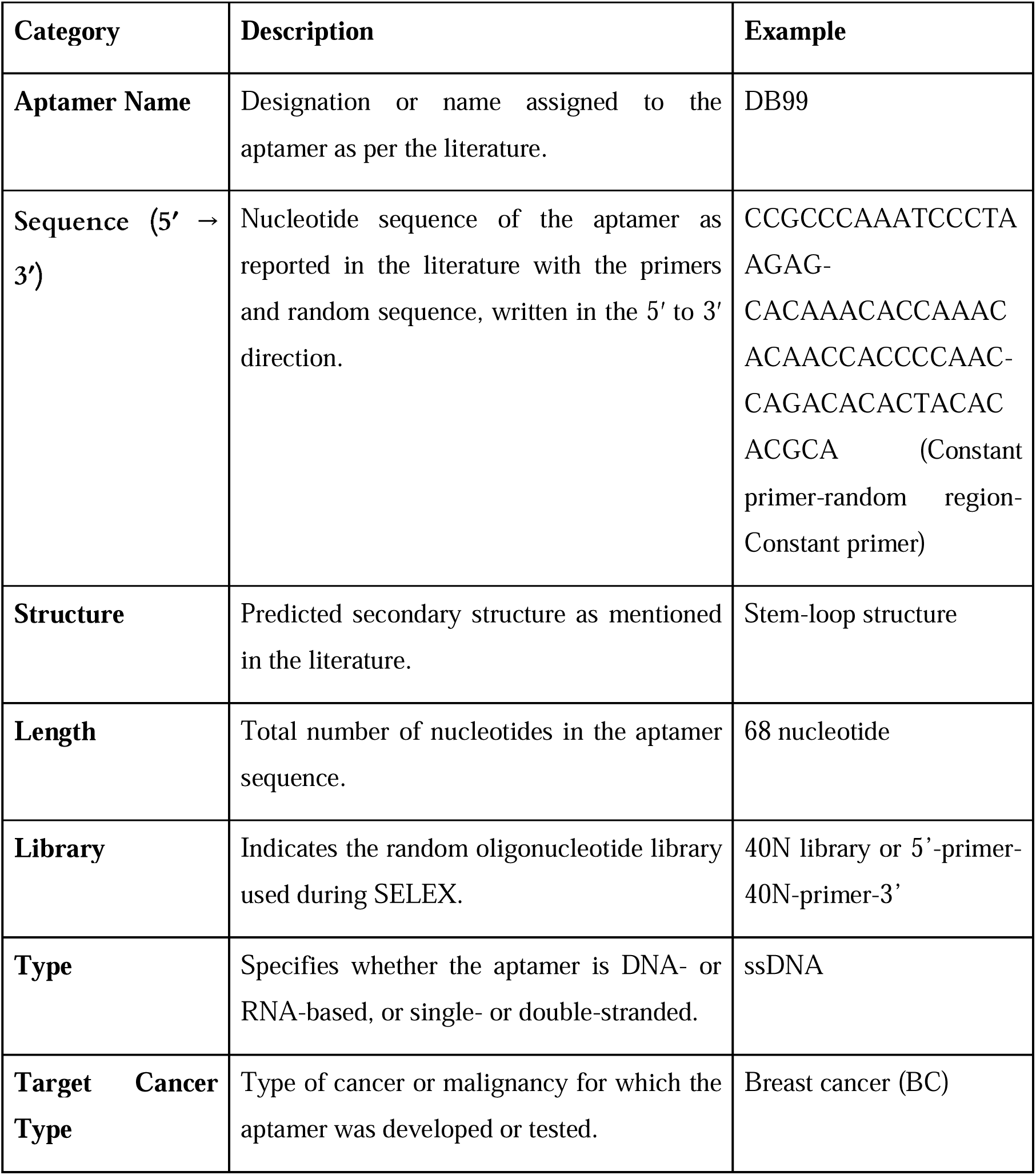
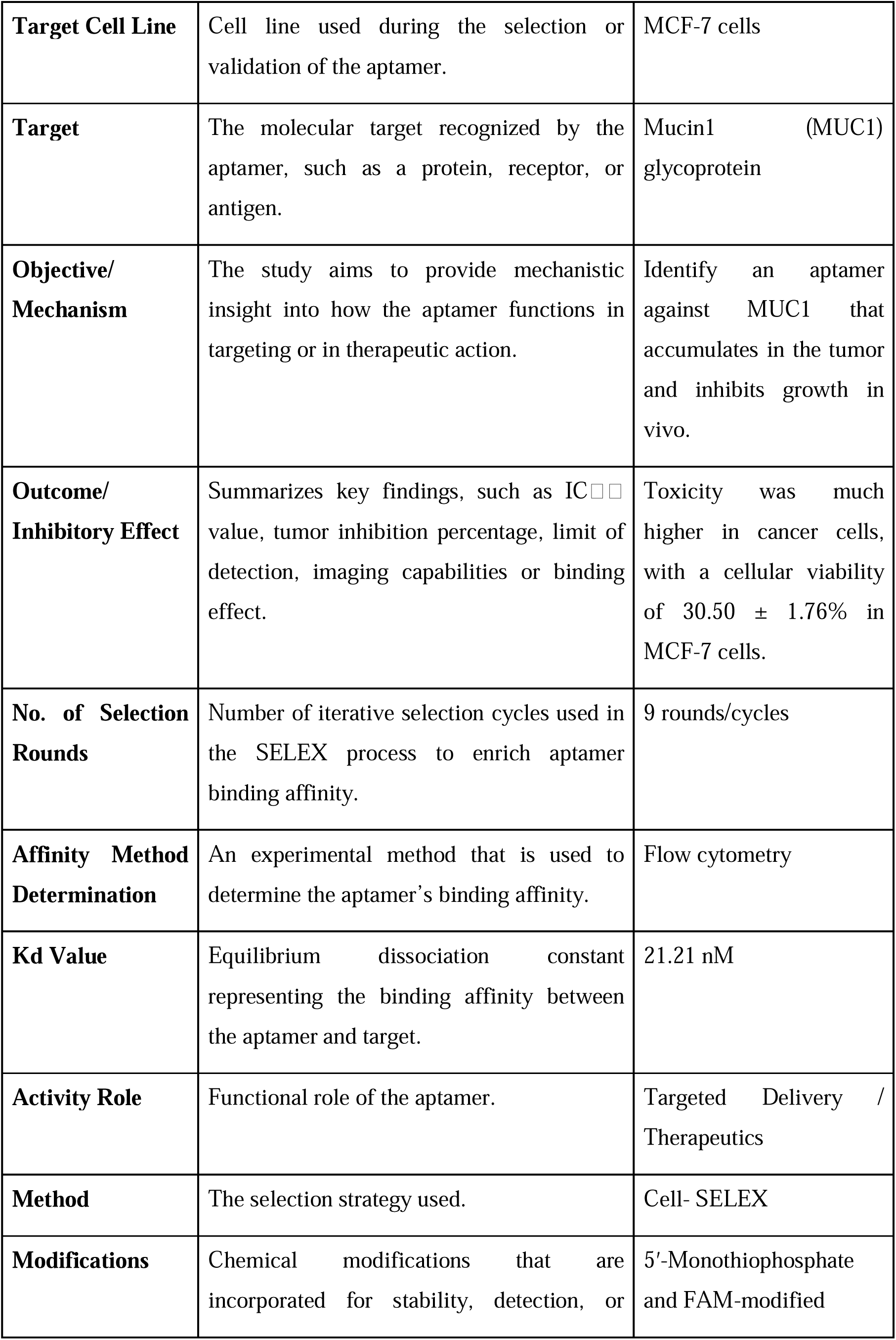
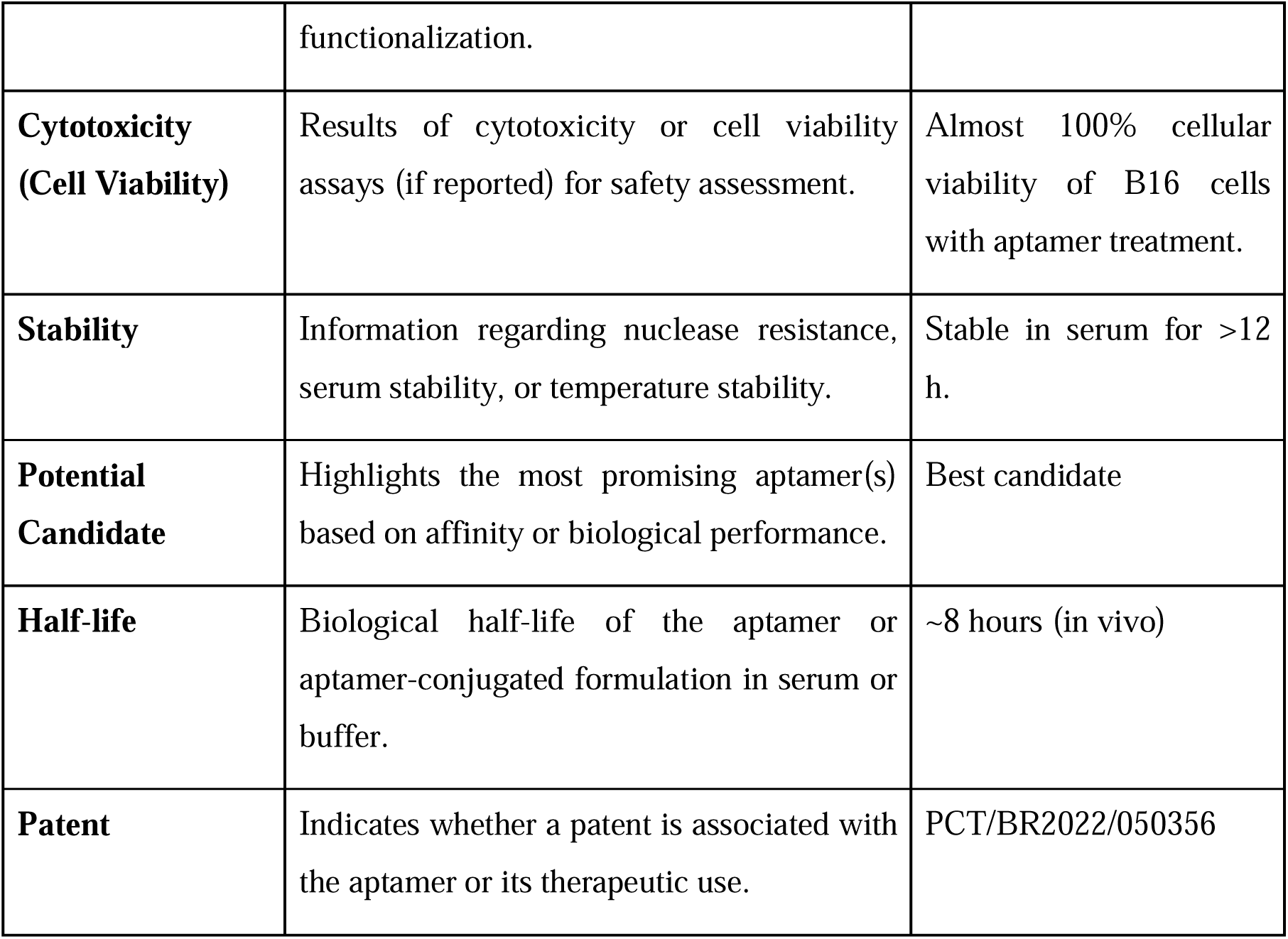
Description of the key fields comprised in the AptCancerDB database.

### 2.3. Functional Overview of Database Modules

#### AptCancerDB Architecture and User Interface

AptCancerDB is built to store and manage data using MySQL (version 5.5.62) and Apache HTTP Server (version 2.4.7). The backend operates PHP scripts to handle database-related requests. The frontend, or user interface, is a web-based and responsive application developed with TypeScript. This application dynamically generates HTML in the browser, making it compatible across platforms. It utilizes React and the Tailwind CSS framework for styling. The interface includes tools for querying, browsing, and analyzing data. Every entry in the system is versioned to keep track of additions and updates. Users can access it for free at https://webs.iiitd.edu.in/raghava/aptcancerdb.

#### Data Querying

Various modules have been developed to facilitate database searches on the web server. A ‘Basic Search’ module forms a key feature that allows for searching based on either the utilizing aptcancer ID or nucleic acid type for running a standard search. The result page will display the requested ID or relevant IDs. The ‘Advanced Search’ option in AptCancerDB allows more sophisticated search queries. It features a multi-query mode, where users can submit multiple requests within a single query or in separate requests, and perform multiple searches using Boolean expressions. The “Secondary Search” feature enables 2D-structure-based aptamer sequence searching. AptCancerDB utilises the dot-bracket notation for representing structures, which is a well-defined type system in which unpaired nucleotides are represented by dots (“.”) and paired bases are indicated by brackets (“()”). It is a popular tool in modelling functional nucleic acids, as it effectively encodes and compares secondary structures (62). AptCancerDB users can search the database based on a query aptamer’s Dot notation, either as a substructure, a superstructure, or an exact fragment.

#### Browsing

For easy access to the database content, the data has been organised by the browsing module. It lists multiple anticancer aptamer-identification submodules for browsing, including ‘Cell Line-wise’, ‘Cancer type-wise’, ‘Length-wise’, ‘Modification-wise’,’ Affinity-wise’, ‘Activity role’, ‘Clinical Trials’, and ‘Publication year’. The database contained approximately 200 cancer cell lines from 29 cancer types. Each entry is linked to the corresponding aptamer card, which contains detailed information about it. Key information on the aptamer card includes the PMID/DOI, sequence, length, origin, and other relevant details. Modifications introduced during the aptamer synthesis process, the affinity methods employed, the cell line used, and the outcomes of preclinical studies are also included. Literature information includes paper title, DOI, journal, and abstract. Structure information includes secondary structure predicted using the ViennaRNA Package 2.0 (63), Dot-notations, structure elements (stem, loop, hairpin, etc.), and the minimum free energy (MFE) value in kcal/mol. We have also included a Cross-Referencing module that displays a table containing all entries in our database, sourced from the other two databases: AptaDB and Aptabase. This module provides links to corresponding resources within these databases for users to access additional information.

#### Analysis Tools

In our databases, users can compare their input nucleotide sequences by performing a similarity search using the Basic Local Alignment Search Tool (BLAST) with the blastn suite, employing closely related anticancer aptamer sequences. This platform enables users to run sub-searches and super-searches using the Mapping. When a sub-search is performed, a query aptamer sequence is mapped against all anticancer-related aptamers present in AptCancerDB to find sequence matches; whereas a super-search returns all aptamers in the database that are similar to the nucleotide sequences given as a query. If no hits are found, then the results page will be empty. Additionally, a GC calculator has been incorporated to count each nucleotide and determine the GC composition of the aptamer sequences, thereby assessing their stability.

#### Knowledge Graph

The knowledge graph was implemented using the ArcadeDB (https://arcadedb.com/knowledge-graphs.html) multi-model graph database, which provides native OpenCypher support. Four vertex types, including Aptamer, Target, CancerType, and ActivityRole, and three directed edge types, including TARGET_EDGE, TARGETS_CANCER_EDGE, and HAS_ACTIVITY_ROLE_EDGE, were defined to represent the biological entities and their interrelationships. Triple files were ingested using OpenCypher LOAD CSV queries with MERGE operations to ensure node deduplication across datasets. This architecture enables multi-hop graph traversal, supporting queries such as identifying aptamers shared across multiple cancer types, mapping target proteins implicated in specific malignancies, and grouping aptamers by their functional roles across cancer subtypes.

#### Download

The complete AptCancerDB data is available for download. For the aptamer sequence, a multifasta file can be downloaded. Also, we allow users to download the predicted secondary structure for the aptamer entries.

#### Rest API

We have added programmatic data retrieval for AptCancerDB in this release to reduce the need for manual downloads. Programs can use simple URLs (REST) to retrieve data by changing the desired parameters. The API returns data in JSON format, a standard data exchange format which can be customized by the users.

## 3. Results and Discussion

### 3.1. Overview of Database Statistics

AptCancerDB compiles 1,941 experimentally validated entries of anticancer aptamers retrieved from 950 PubMed-indexed studies and 36 DOI sources spanning 2001-2025, providing a unified resource that integrates sequence, target, cancer type, and functional activity data. As represented in Figure 2(A), 82.7% of entries are categorizes into ssDNA reflecting their higher stability, ease of synthesis, and frequent use in cell-SELEX. and 16.6% of ssRNA, especially those selected against intracellular or protein-protein interaction targets, where structural flexibility is advantageous. Importantly, DNA aptamers exhibit inherently greater stability than RNA aptamers, largely due to the absence of the 2_′_-hydroxyl group in DNA sugars, which reduces chemical reactivity and susceptibility to nuclease degradation. Aldo structural differences, such as variations in helix form, can make DNA more resistant to nucleases (64). For these reasons, DNA aptamers are often preferred in therapeutic applications and biomedical research (65).

**Figure 2:**
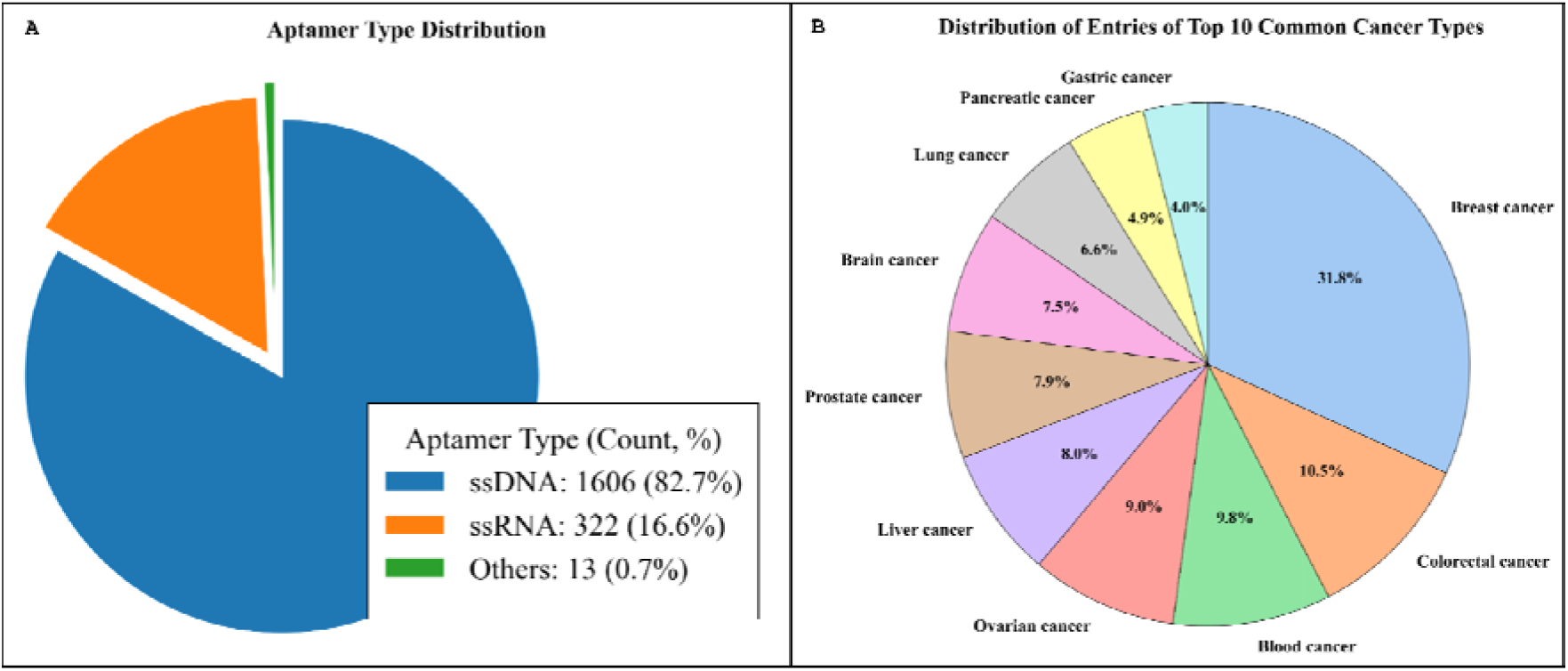
Pie charts showing (A) Distribution of aptamers based on the type of nucleic acids. (B) The percentage of entries from the top 10 cancer types.

The database includes entries from 29 different types of cancers, with breast cancer showing the highest representation (31.8%), followed by colorectal, blood, ovarian, and liver cancers. As depicted in Figure 2(B), these categories account for over two-thirds of all entries. Other cancers, such as prostate, brain, lung, and pancreatic, showed intermediate indications, reflecting rising interest in hard-to-treat tumors. Consistent with these patterns, the selection and validation of aptamers depend on some widely used cancer cell lines. As shown in Figure 3(A), MCF-7 and MDA-MB-231, both breast adenocarcinoma cell lines, emerged as the most frequently used models, followed by SK-BR-3, HepG2, and LNCaP. Figure 3(B) represents the commonly used targets for aptamer discovery; among them, the dominance of whole-cell targets (405 entries) and tumor-associated markers or receptors such as nucleolin (NCL), MUC1 glycoprotein, PTK7, PSMA, EpCAM, EGFR, and HER2 highlights their importance in cancer biology. These commonly targeted proteins and cell-surface molecules anchor the majority of therapeutic and diagnostic strategies explored in current aptamer research.

**Figure 3:**
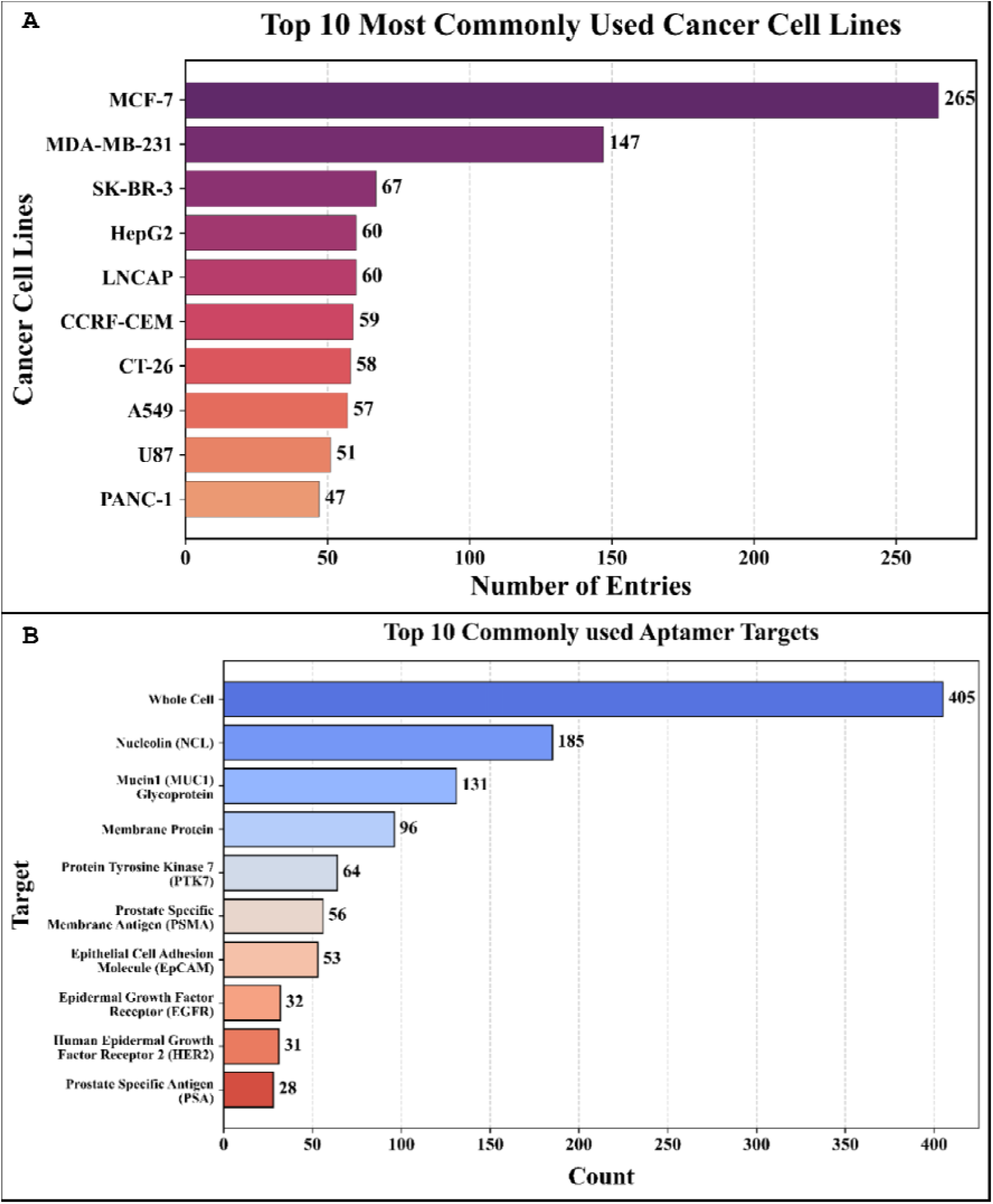
Bar plots showing (A) Top 10 most common cancer cell lines, and (B) Top 10 most commonly used aptamer targets for cancer studies.

Likewise, the cumulative year-wise growth pattern shown in Figure 4 indicates a rapid increase in anticancer aptamer publications, particularly after 2013, despite the first aptamer drug, Pegaptanib (Macugen), being approved by the FDA in 2004. The number of entries increased from 346 between 2009 and 2012 to 924 from 2013 to 2016, and further to 1,517 (2017-2020), and ultimately reached 1,941 after 2025. This steady rise indicates growing interest in aptamer-based diagnostics, drug delivery, imaging, and therapeutics, driven by advancements in SELEX technologies, next-generation sequencing (NGS), nanomedicine, and computational methods.

**Figure 4:**
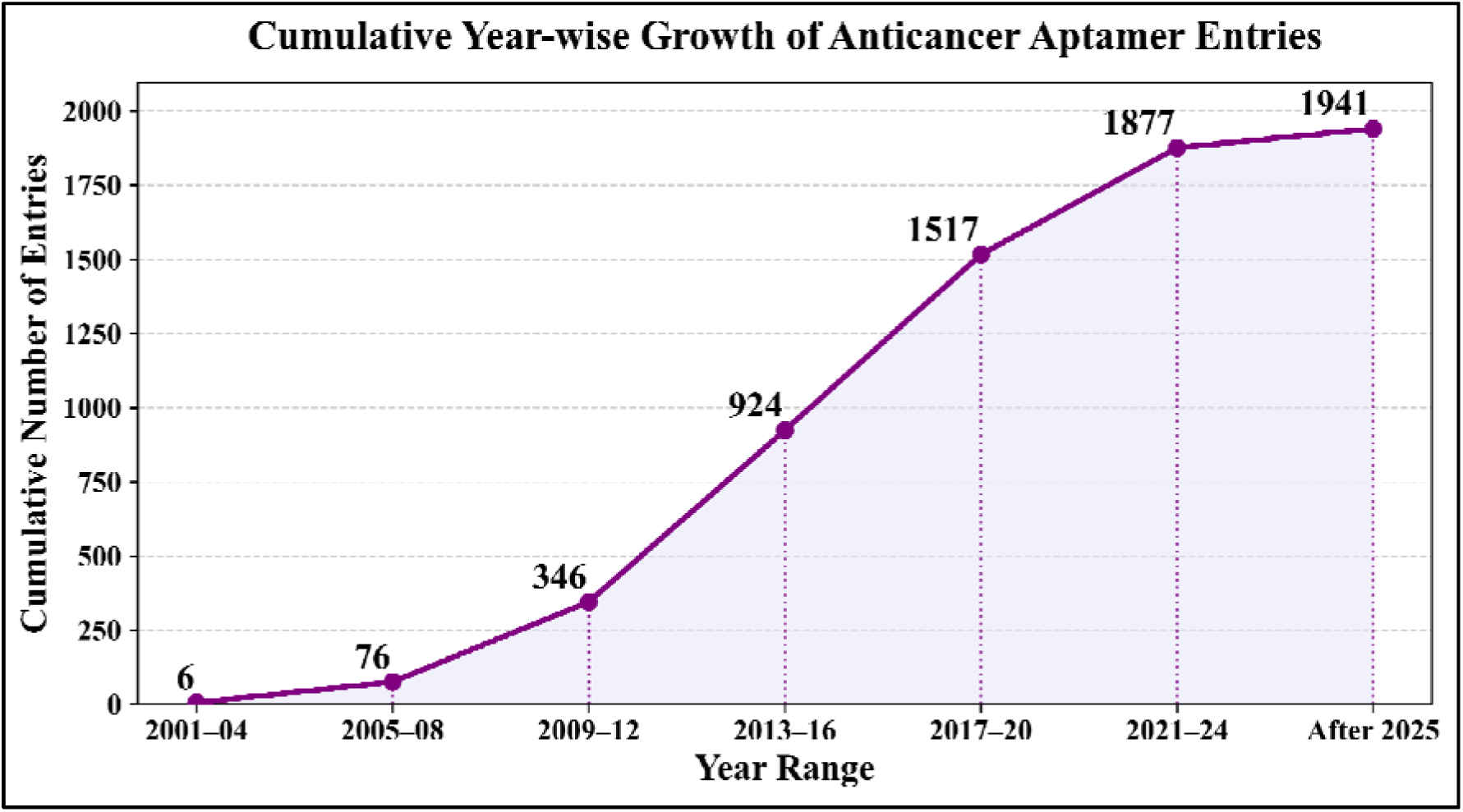
Line plot showing the growth trend of aptamers in the field of cancer studies.

Similarly in the database, aptamers exhibit broad versatility in their functional roles, encompassing therapeutic, inhibitory, diagnostic, detection, imaging, delivery, immunotherapy, and biosensor applications. As shown in Figure 5 (A), aptamer sequence lengths exhibit a characteristic trend, with most functional aptamers typically found in the 70-90 nucleotide (nt) range, which helps them maintain stable secondary and tertiary structures essential for recognizing their targets. Shorter aptamers, typically around 30-50 nt, often represent truncated, affinity-optimised derivatives produced through iterative structure-activity refinement. On the other hand, very long aptamers, those exceeding 111 nt, are uncommon, likely due to high synthetic costs, structural redundancy, and reduced stability. Figure 5 (A) also reveals distinct length preferences across various activity roles, for example, therapeutic and detection aptamers dominate the 70-90 nt range, which plays important role in blocking oncogenic proteins or receptors for downstream signaling, whereas differences lie as a significant number are also involved in diagnostic, imaging and detection-based roles, including biosensors and imaging probes for the identification of whole cancer cells or the quantification of tumor-biomarkers. Another major group focuses on targeted delivery (524 entries), where aptamers guide drugs, genes, mRNA, siRNAs, liposomes or nanoparticles selectively to tumor sites for minimizing the systemic toxicity. This reflects the diversity of selection strategies, target types, and structural requirements underlying each biological function. Although smaller in number, entries in immunotherapy emphasize aptamers’ growing integration into combined therapeutic systems involving immunocomplex reactions.

**Figure 5:**
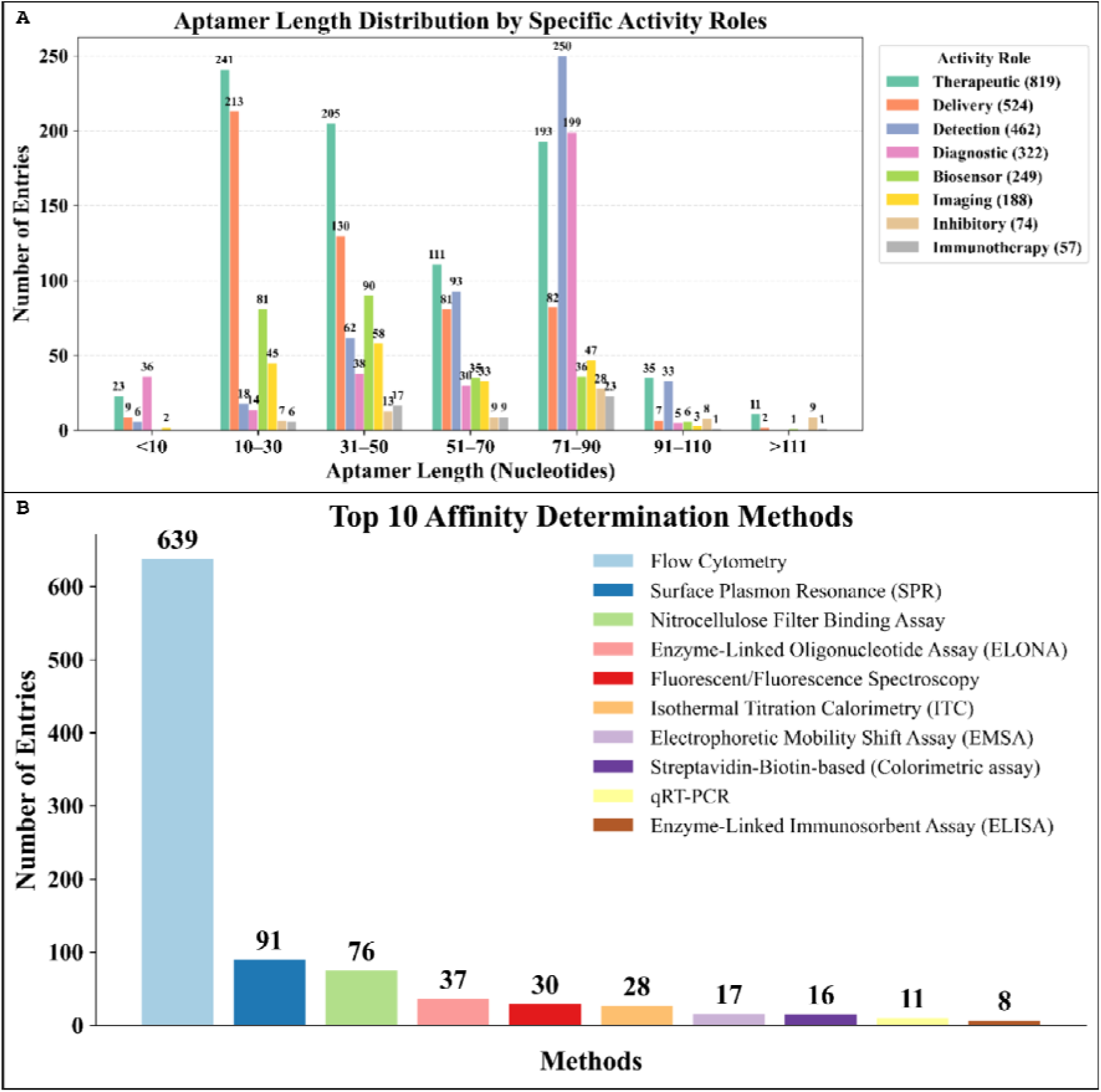
(A) Distribution of different activity roles according to the length of the aptamer. (B) Bar plot showing the top 10 most commonly used methods for affinity determination.

AptCancerDB also features the experimental methods used to assess aptamer binding affinities. Figure 5 (B) summarizes the top affinity-determination approaches, showing Flow Cytometry as the most dominant assay (639 entries), which is followed by Surface Plasmon Resonance (SPR), Nitrocellulose Filter Binding, ELONA, and various fluorescence-based techniques. These patterns indicate a growing emphasis on quantitative, high-sensitivity binding assays, which are beneficial for translational and preclinical research. The wide range of methods illustrates the diversity of aptamer applications, from targeting cells to exploring protein-aptamer interactions.

### 3.2. Secondary Structure and Thermodynamic Stability Analysis

Knowing the 2D secondary structures of aptamers is essential not only for understanding folding behaviour but also for serving as a critical input for 3D structural modelling and downstream computational design workflows (66). Therefore, to assess structural robustness, secondary structures were predicted, and MFEs were calculated for all curated aptamer sequences. The MFE values provide an estimate of thermodynamic stability, with lower (more negative) values indicating more energetically favourable folding; hence, aptamers with very low MFE values may be promising candidates for high-specificity binding or 3D modelling. In Figure 6(A), the distribution of MFE values exhibits a distinct left skew, with an average of approximately −6.35 kcal/mol, indicating that most aptamers fold into thermodynamically stable secondary structures, which are essential for high-affinity binding to their targets. Similarly, in Figure 6(B), a moderated negative correlation between sequence length and MFE was also observed, suggesting that longer sequences form more complex and energetically favourable structures which confirms why many of the curated functional aptamers fall into the ∼40–90 nucleotide range (in Figure 5 (A)) which are long enough to form necessary structures but not so long as to increase synthesis cost or instability (67). This trend aligns with expected nucleic-acid folding behaviour, in which increased nucleotide content enables the formation of additional stems, loops, and tertiary motifs, thereby lowering the free energy.

**Figure 6:**
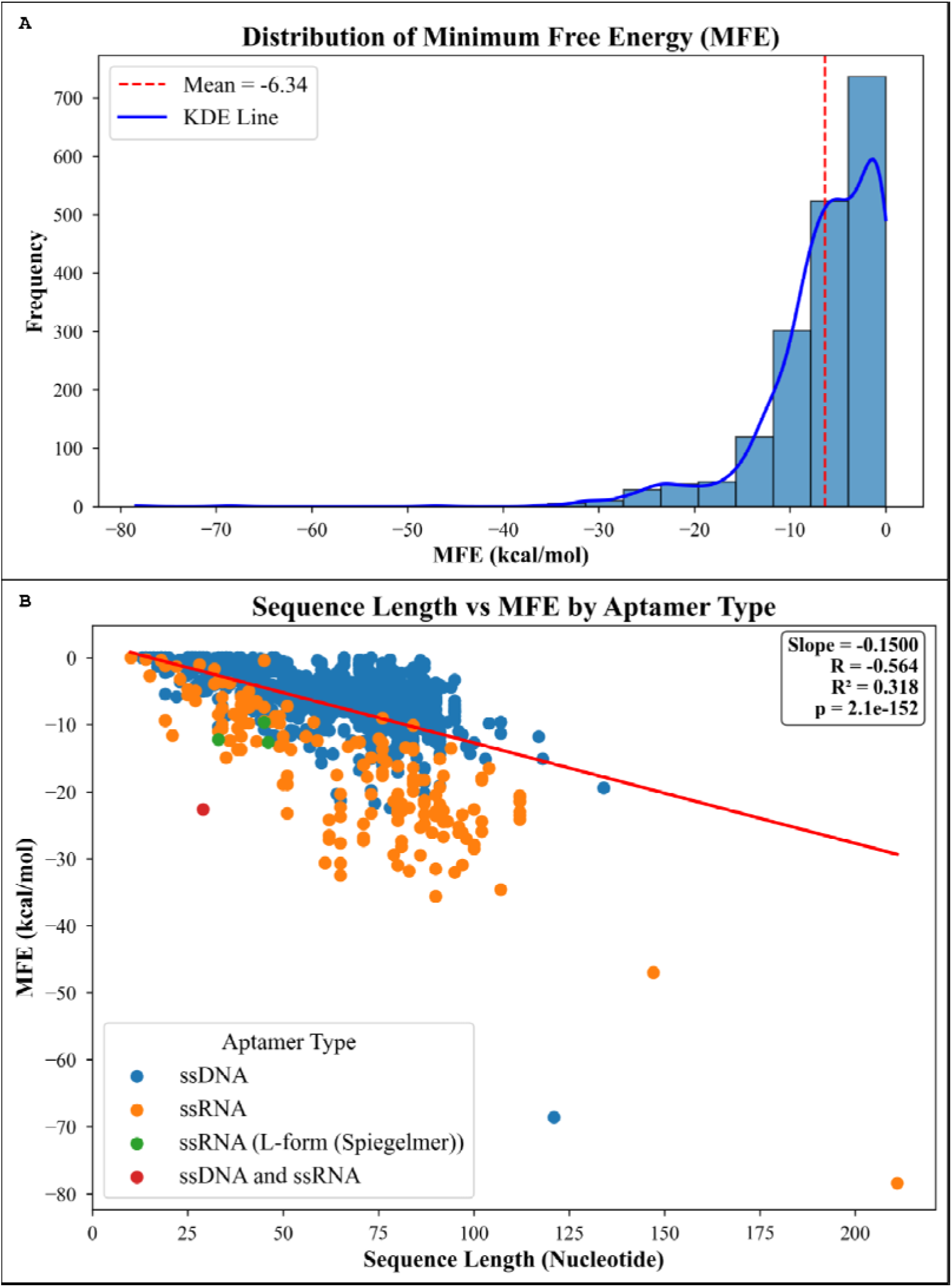
(A)Bar plot showing the distribution of predicted MFE values across the dataset. (B) Regression plot of aptamer sequence length vs predicted MFE.

These results highlight the stability of cancer-targeting aptamers and demonstrate how aptamer research has progressed from discovery to practical application. In general, this resource focuses on cancer, helps identify, rationally design, and compare various aptamer candidates.

## 4. Future Prospects

The future of cancer aptamer research will be guided by computational design, the identification of new targets, and studies that translate lab-based ideas into real-world applications. Curated resources like AptCancerDB, along with machine learning and motif analysis, help to enhance aptamer-target prediction, leading to designing tumor-homing aptamers that can enhance penetration into tumors, retention, and specificity. Expanding the focus to include markers of drug resistance, factors in the tumor environment, and immune checkpoint ligands could uncover new treatment alternatives. Also, combining aptamer-based drug or gene delivery with immunotherapies shows great promise for treating cancer. In the coming years, we plan to enhance the database by incorporating patent information, thereby transforming AptCancerDB into a dynamic resource that facilitates the development of next-generation aptamers for cancer treatments.

## 5. Conclusion

In summary, AptCancerDB is a platform that includes 1941 aptamers specifically related to cancer. It provides tools for searching, browsing, and analyzing the data, which helps in comparing different aptamers and designing more effective ones. By consolidating information from various sources into a focused cancer resource, AptCancerDB facilitates research in the development of new cancer treatments. Users can access the data by downloading it directly from the website or using programming methods. By integrating and standardising this information, we aim to support researchers in both the computational design of aptamers and the translational development of aptamer-based therapies and diagnostics in cancer research. As the field rapidly evolves toward nanocarrier systems, stimuli-responsive delivery, and immunotherapeutic applications, aptamers are poised to play an increasingly prominent role in precision oncology. We invite the community to explore the database, contribute new entries, and utilise the resource to accelerate aptamer-based innovations in cancer therapy and diagnostics.

## Acknowledgements

The authors are grateful to the Council of Scientific and Industrial Research (CSIR), the Department of Biotechnology (DBT), and the Department of Computational Biology at IIITD, New Delhi, for providing fellowships and financial support, as well as for the necessary infrastructure and facilities. We thank Draw.io for creating the Figures.

## Author Contributions

The data were collected, curated and analyzed by NB. The web server’s front-end and back-end were developed by SS. The manuscript was prepared by NB, SS, and GPSR. GPSR conceived the idea and coordinated the project. All writers have reviewed and approved the final version of the paper.

## Data availability

We will update it soon.

## Declaration of generative AI and AI-assisted technologies

During the preparation of this work, the author(s) used ChatGPT in order to improve the writing and language of the manuscript. After using this tool/service, the author(s) reviewed and edited the content as needed and take(s) full responsibility for the content of the published article.

## References

1. Pereira, R.L., Nascimento, I.C., Santos, A.P., Ogusuku, I.E.Y., Lameu, C., Mayer, G. and Ulrich, H. (2018) Aptamers: novelty tools for cancer biology. Oncotarget, 9, 26934–26953.

2. Siegel, R.L., Kratzer, T.B., Giaquinto, A.N., Sung, H. and Jemal, A. (2025) Cancer statistics, 2025. CA Cancer J Clin, 75, 10–45.

3. Karpuz, M., Silindir-Gunay, M. and Ozer, A.Y. (2018) Current and Future Approaches for Effective Cancer Imaging and Treatment. Cancer Biother Radiopharm, 33, 39–51.

4. Li, Y., Zamay, T.N., Luzan, N.A., Pryakhin, E.A., Styazhkina, E.V., Osminkina, L.A., Kolovskaya, O.S., Dymova, M.A., Kuligina, E.V., Richter, V.A., et al. (2025) Aptamers as a new frontier in targeted cancer therapy. Adv Drug Deliv Rev, 226, 115692.

5. May, M. (2022) Why drug delivery is the key to new medicines. Nat Med, 28, 1100–1102.

6. Khedri, M., Rafatpanah, H., Abnous, K., Ramezani, P. and Ramezani, M. (2015) Cancer immunotherapy via nucleic acid aptamers. Int Immunopharmacol, 29, 926–936.

7. Mahmoudian, F., Ahmari, A., Shabani, S., Sadeghi, B., Fahimirad, S. and Fattahi, F. (2024) Aptamers as an approach to targeted cancer therapy. Cancer Cell Int, 24, 108.

8. Zhu, G. and Chen, X. (2018) Aptamer-based targeted therapy. Adv Drug Deliv Rev, 134, 65–78.

9. Mayer, G. (2009) The chemical biology of aptamers. Angew Chem Int Ed Engl, 48, 2672–2689.

10. Ku, T.-H., Zhang, T., Luo, H., Yen, T.M., Chen, P.-W., Han, Y. and Lo, Y.-H. (2015) Nucleic Acid Aptamers: An Emerging Tool for Biotechnology and Biomedical Sensing. Sensors (Basel), 15, 16281–16313.

11. Tuerk, C. and Gold, L. (1990) Systematic evolution of ligands by exponential enrichment: RNA ligands to bacteriophage T4 DNA polymerase. Science, 249, 505–510.

12. Cesarini, V., Appleton, S.L., de Franciscis, V. and Catalucci, D. (2025) The recent blooming of therapeutic aptamers. Mol Aspects Med, 102, 101350.

13. Fang, Z., Feng, X., Tang, F., Jiang, H., Han, S., Tao, R. and Lu, C. (2024) Aptamer Screening: Current Methods and Future Trend towards Non-SELEX Approach. Biosensors (Basel), 14.

14. Mili, M., Bachu, V., Kuri, P.R., Singh, N.K. and Goswami, P. (2024) Improving synthesis and binding affinities of nucleic acid aptamers and their therapeutics and diagnostic applications. Biophys Chem, 309, 107218.

15. Kohlberger, M. and Gadermaier, G. (2022) SELEX: Critical factors and optimization strategies for successful aptamer selection. Biotechnol Appl Biochem, 69, 1771–1792.

16. Gold, L. (1995) Oligonucleotides as research, diagnostic, and therapeutic agents. J Biol Chem, 270, 13581–13584.

17. Dassie, J.P., Liu, X.-Y., Thomas, G.S., Whitaker, R.M., Thiel, K.W., Stockdale, K.R., Meyerholz, D.K., McCaffrey, A.P., McNamara, J.O., 2nd and Giangrande, P.H. (2009) Systemic administration of optimized aptamer-siRNA chimeras promotes regression of PSMA-expressing tumors. Nat Biotechnol, 27, 839–849.

18. Zhang, Y., Hong, H. and Cai, W. (2011) Tumor-targeted drug delivery with aptamers. Curr Med Chem, 18, 4185–4194.

19. Steurer, M., Montillo, M., Scarfò, L., Mauro, F.R., Andel, J., Wildner, S., Trentin, L., Janssens, A., Burgstaller, S., Frömming, A., et al. (2019) Olaptesed pegol (NOX-A12) with bendamustine and rituximab: a phase IIa study in patients with relapsed/refractory chronic lymphocytic leukemia. Haematologica, 104, 2053–2060.

20. Yang, C., Zhao, H., Sun, Y., Wang, C., Geng, X., Wang, R., Tang, L., Han, D., Liu, J. and Tan, W. (2022) Programmable manipulation of oligonucleotide-albumin interaction for elongated circulation time. Nucleic Acids Res, 50, 3083–3095.

21. Liu, S., Li, X., Gao, H., Chen, J. and Jiang, H. (2025) Progress in Aptamer Research and Future Applications. ChemistryOpen, 14, e202400463.

22. Mahlknecht, G., Maron, R., Mancini, M., Schechter, B., Sela, M. and Yarden, Y. (2013) Aptamer to ErbB-2/HER2 enhances degradation of the target and inhibits tumorigenic growth. Proc Natl Acad Sci U S A, 110, 8170–8175.

23. Hoellenriegel, J., Zboralski, D., Maasch, C., Rosin, N.Y., Wierda, W.G., Keating, M.J., Kruschinski, A. and Burger, J.A. (2014) The Spiegelmer NOX-A12, a novel CXCL12 inhibitor, interferes with chronic lymphocytic leukemia cell motility and causes chemosensitization. Blood, 123, 1032–1039.

24. Camorani, S., Crescenzi, E., Colecchia, D., Carpentieri, A., Amoresano, A., Fedele, M., Chiariello, M. and Cerchia, L. (2015) Aptamer targeting EGFRvIII mutant hampers its constitutive autophosphorylation and affects migration, invasion and proliferation of glioblastoma cells. Oncotarget, 6, 37570–37587.

25. Chinnadurai, R.K., Khan, N., Meghwanshi, G.K., Ponne, S., Althobiti, M. and Kumar, R. (2023) Current research status of anti-cancer peptides: Mechanism of action, production, and clinical applications. Biomed Pharmacother, 164, 114996.

26. Gautam, A., Kapoor, P., Chaudhary, K., Kumar, R., Open Source Drug Discovery Consortium and Raghava, G.P.S. (2014) Tumor homing peptides as molecular probes for cancer therapeutics, diagnostics and theranostics. Curr Med Chem, 21, 2367–2391.

27. Doherty, C.D., Wilbanks, B.A., Jain, S., S Pearson, K., Bakken, K.K., Burgenske, D.M., Lett, N.W., Sarkaria, J.N. and Maher, L.J., 3rd (2025) selection of anti-glioblastoma DNA aptamers in an orthotopic patient-derived xenograft model. NAR Cancer, 7, zcaf005.

28. Kim, D.-H., Seo, J.-M., Shin, K.-J. and Yang, S.-G. (2021) Design and clinical developments of aptamer-drug conjugates for targeted cancer therapy. Biomater Res, 25, 42.

29. Sanati, M., Afshari, A.R., Ahmadi, S.S., Kesharwani, P. and Sahebkar, A. (2023) Aptamers against cancer drug resistance: Small fighters switching tactics in the face of defeat. Biochim Biophys Acta Mol Basis Dis, 1869, 166720.

30. Gautam, A., Chaudhary, K., Kumar, R. and Raghava, G.P.S. (2015) Computer-Aided Virtual Screening and Designing of Cell-Penetrating Peptides. Methods Mol Biol, 1324, 59–69.

31. Dzarieva, F.M., Pavlova, S.A., Pavlova, G.V. and Fab, L.V. (2025) Aptamers and endocytosis: The path into tumor cells. Biochem Biophys Res Commun, 791, 152960.

32. Lee, J.W., Kim, H.J. and Heo, K. (2015) Therapeutic aptamers: developmental potential as anticancer drugs. BMB Rep, 48, 234–237.

33. Zhou, J. and Rossi, J. (2017) Aptamers as targeted therapeutics: current potential and challenges. Nat Rev Drug Discov, 16, 181–202.

34. Morita, Y., Leslie, M., Kameyama, H., Volk, D.E. and Tanaka, T. (2018) Aptamer Therapeutics in Cancer: Current and Future. Cancers (Basel), 10.

35. Shaban, S.M. and Kim, D.-H. (2021) Recent Advances in Aptamer Sensors. Sensors (Basel), 21.

36. Sequeira-Antunes, B. and Ferreira, H.A. (2023) Nucleic Acid Aptamer-Based Biosensors: A Review. Biomedicines, 11.

37. Yang, W., Ran, C., Lian, X., Wang, Z., Du, Z., Bing, T., Zhang, Y. and Tan, W. (2025) Aptamer-based targeted drug delivery and disease therapy in preclinical and clinical applications. Adv Drug Deliv Rev, 226, 115680.

38. Han, J., Gao, L., Wang, J. and Wang, J. (2020) Application and development of aptamer in cancer: from clinical diagnosis to cancer therapy. J Cancer, 11, 6902–6915.

39. Gao, F., Yin, J., Chen, Y., Guo, C., Hu, H. and Su, J. (2022) Recent advances in aptamer-based targeted drug delivery systems for cancer therapy. Front Bioeng Biotechnol, 10, 972933.

40. Agnello, L., d’Argenio, A., Nilo, R., Fedele, M., Camorani, S. and Cerchia, L. (2023) Aptamer-Based Strategies to Boost Immunotherapy in TNBC. Cancers (Basel), 15.

41. Anantha Rajah, D., Tan, H.S. and Farghadani, R. (2025) Aptamers as immune checkpoint inhibitors in cancer immunotherapy: targeting CTLA-4/B7 and PD-1/PD-L1 pathways. Int Immunopharmacol, 164, 115339.

42. Stoltenburg, R., Reinemann, C. and Strehlitz, B. (2007) SELEX--a (r)evolutionary method to generate high-affinity nucleic acid ligands. Biomol Eng, 24, 381–403.

43. Bashir, A., Yang, Q., Wang, J., Hoyer, S., Chou, W., McLean, C., Davis, G., Gong, Q., Armstrong, Z., Jang, J., et al. (2021) Machine learning guided aptamer refinement and discovery. Nat Commun, 12, 2366.

44. Chen, Z., Hu, L., Zhang, B.-T., Lu, A., Wang, Y., Yu, Y. and Zhang, G. (2021) Artificial Intelligence in Aptamer-Target Binding Prediction. Int J Mol Sci, 22.

45. Lee, J.F., Hesselberth, J.R., Meyers, L.A. and Ellington, A.D. (2004) Aptamer database. Nucleic Acids Res, 32, D95–100.

46. Askari, A., Kota, S., Ferrell, H., Swamy, S., Goodman, K.S., Okoro, C.C., Spruell Crenshaw, I.C., Hernandez, D.K., Oliphant, T.E., Badrayani, A.A., et al. (2024) UTexas Aptamer Database: the collection and long-term preservation of aptamer sequence information. Nucleic Acids Res, 52, D351–D359.

47. Chen, L., Yu, Z., Wu, Z., Zhou, M., Wang, Y., Yu, X., Li, W., Liu, G. and Tang, Y. (2024) AptaDB: a comprehensive database integrating aptamer-target interactions. RNA, 30, 189–199.

48. Ponomarenko, J.V., Orlova, G.V., Frolov, A.S., Gelfand, M.S. and Ponomarenko, M.P. (2002) SELEX_DB: a database on in vitro selected oligomers adapted for recognizing natural sites and for analyzing both SNPs and site-directed mutagenesis data. Nucleic Acids Res, 30, 195–199.

49. Thodima, V., Pirooznia, M. and Deng, Y. (2006) RiboaptDB: a comprehensive database of ribozymes and aptamers. BMC Bioinformatics, 7 Suppl 2, S6.

50. Cruz-Toledo, J., McKeague, M., Zhang, X., Giamberardino, A., McConnell, E., Francis, T., DeRosa, M.C. and Dumontier, M. (2012) Aptamer Base: a collaborative knowledge base to describe aptamers and SELEX experiments. Database (Oxford), 2012, bas006.

51. Lu, Z., Sun, H., Fu, B., Ao, Y., Wang, J., Chen, K., Luo, Y., Li, L., Qiu, Z., Zhao, J., et al. (2026) Ribocentre-aptamer: an integrative, structure-focused database for RNA aptamers. Nucleic Acids Res, 54, D264–D272.

52. Kumar, R., Chaudhary, K., Gupta, S., Singh, H., Kumar, S., Gautam, A., Kapoor, P. and Raghava, G.P.S. (2013) CancerDR: cancer drug resistance database. Sci Rep, 3, 1445.

53. Faraji, N., Arab, S.S., Doustmohammadi, A., Daly, N.L. and Khosroushahi, A.Y. (2022) ApInAPDB: a database of apoptosis-inducing anticancer peptides. Sci Rep, 12, 21341.

54. Chauhan, M., Gupta, A., Tomer, R. and Raghava, G.P.S. (2025) CancerPPD2: an updated repository of anticancer peptides and proteins. Database (Oxford), 2025.

55. Wang, G., Schmidt, C., Li, X. and Wang, Z. (2026) APD6: the antimicrobial peptide database is expanded to promote research and development by deploying an unprecedented information pipeline. Nucleic Acids Res, 54, D363–D374.

56. Gupta, S., Chaudhary, K., Kumar, R., Gautam, A., Nanda, J.S., Dhanda, S.K., Brahmachari, S.K. and Raghava, G.P.S. (2016) Prioritization of anticancer drugs against a cancer using genomic features of cancer cells: A step towards personalized medicine. Sci Rep, 6, 23857.

57. Agrawal, P., Bhagat, D., Mahalwal, M., Sharma, N. and Raghava, G.P.S. (2021) AntiCP 2.0: an updated model for predicting anticancer peptides. Brief Bioinform, 22.

58. Arif, M., Musleh, S., Fida, H. and Alam, T. (2024) PLMACPred prediction of anticancer peptides based on protein language model and wavelet denoising transformation. Sci Rep, 14, 16992.

59. Gao, S., Xia, Y., Li, X., Cui, F., Zhang, Q., Zou, Q. and Zhang, Z. (2025) ACP-ESM2: Enhancing Anticancer Peptide Prediction With Pre-Trained Protein Language Models. IEEE Trans Comput Biol Bioinform, 22, 1041–1051.

60. Li, Z., Fu, X., Huang, J., Zeng, P., Huang, Y., Chen, X. and Liang, C. (2021) Advances in Screening and Development of Therapeutic Aptamers Against Cancer Cells. Front Cell Dev Biol, 9, 662791.

61. Shigdar, S., Schrand, B., Giangrande, P.H. and de Franciscis, V. (2021) Aptamers: Cutting edge of cancer therapies. Mol Ther, 29, 2396–2411.

62. Ramlan, E.I. and Zauner, K.-P. (2013) In-silico design of computational nucleic acids for molecular information processing. J Cheminform, 5, 22.

63. Lorenz, R., Bernhart, S.H., Höner Zu Siederdissen, C., Tafer, H., Flamm, C., Stadler, P.F. and Hofacker, I.L. (2011) ViennaRNA Package 2.0. Algorithms Mol Biol, 6, 26.

64. Kratschmer, C. and Levy, M. (2017) Effect of Chemical Modifications on Aptamer Stability in Serum. Nucleic Acid Ther, 27, 335–344.

65. Afanasyeva, A., Nagao, C. and Mizuguchi, K. (2019) Prediction of the secondary structure of short DNA aptamers. Biophys Physicobiol, 16, 287–294.

66. Popenda, M., Szachniuk, M., Antczak, M., Purzycka, K.J., Lukasiak, P., Bartol, N., Blazewicz, J. and Adamiak, R.W. (2012) Automated 3D structure composition for large RNAs. Nucleic Acids Res, 40, e112.

67. Mokgopa, K.P., Oloniiju, S.D., Lobb, K.A. and Tshiwawa, T. (2025) Benchmarking the Base Randomization Algorithm as a Possible Tool for the Initial Step of Generating a Virtual RNA Aptamers Library. BioTech (Basel), 14.

